# Dynamics of genetic variation in Transcription Factors and its implications for the evolution of regulatory networks in Bacteria

**DOI:** 10.1101/785691

**Authors:** Farhan Ali, Aswin Sai Narain Seshasayee

**Affiliations:** National Centre for Biological Sciences, Tata Institute of Fundamental Research, Bengaluru, Karnataka, 560065, India; Manipal Academy of Higher Education, Manipal, Karnataka, 576104, India

## Abstract

The evolution of bacterial regulatory networks has largely been explained at macroevolutionary scales through lateral gene transfer and gene duplication. Transcription factors (TF) have been found to be less conserved across species than their target genes (TG). This would be expected if TFs accumulate mutations faster than TGs. This hypothesis is supported by several lab evolution studies which found TFs, especially global regulators, to be frequently mutated. Despite these studies, the contribution of point mutations in TFs to the evolution of regulatory network is poorly understood. We tested if TFs show greater genetic variation than their TGs using whole-genome sequencing data from a large collection of *E coli* isolates. We found TFs to be less diverse, across natural isolates, due to their regulatory roles. TFs were enriched in mutations in multiple adaptive lab evolution studies but not in mutation accumulation. However, over long-term evolution, relative frequency of mutations in TFs showed a gradual decay after a rapid initial burst. Our results suggest that point mutations, conferring large-scale expression changes, may drive the early stages of adaptation but gene regulation is subjected to stronger purifying selection post adaptation.

## Introduction

The dynamic environments colonized by bacteria demand optimal regulation of gene expression (1). Transcription initiation, the primary checkpoint in this regulation, is influenced by the activity of a set of DNA-binding proteins called Transcription Factors (TF). TFs sense the cellular environment and respond by activating or suppressing the expression of their target genes (TG). Different species of bacteria occupying diverse niches, thus differ more in the set of their TFs than that of their TGs (2).

The set of transcriptional regulatory interactions in an organism is usually represented as a transcriptional regulatory network (TRN). TRNs have been found to evolve faster than other biological networks (3), based on detection of orthologs across species. Their evolution has been explained largely by duplication (4) and horizontal gene transfer (HGT) (5). Even though both of these processes are accompanied/followed by DNA sequence level changes in the TFs (6, 7), the contribution of point mutations to TRN evolution is poorly understood (8). The significance of point mutations can be realized by the fact that, even where both a TF and its TG are present, the regulatory interaction is often not conserved (9).

Macroevolutionary changes in TRN can be explained, in principle, through mutations at microevolutionary scales, *i.e.*, mutations may accumulate faster in TFs than in TGs within species, and this would be reflected as lower conservation of TFs across species. If selection drives TFs evolution, populations adapting to different environments may select for different mutations in TFs at a higher frequency than in TGs. Some of these may be loss-of-function mutations, leading to complete loss of the TF over a long period of time. In contrast, if TFs evolve through neutral processes, a population may have more standing genetic variation in TFs than in TGs, presumably due to weaker selective constraints. For the same reason, the loss of a TF at large evolutionary distance would be more likely than that of a TG.

The adaptive evolution hypothesis seems to be supported by multiple lab evolution experiments, which found many beneficial mutations to occur in TFs (10). However, this cannot be concluded in the absence of a statistical analysis, of enrichment of beneficial mutations in TFs over TGs, across multiple such studies. Often, these mutations were found in the hubs of the TRN (11), generally referred to as “global” regulators (GR). As TRN follows a power-law distribution of edges per nodes (12), only a few TFs act as GRs, and influence gene expression on a global scale. In this regard, majority of the TFs are considered as “fine-tuners”, and it is not evident if mutations in these TFs should also be more adaptive than in TGs.

The role of adaptation in shaping gene regulation across wild strains also has been demonstrated by several studies (13–15). However, it’s still not clear to what extent mutations in TFs drive regulatory diversification. Some studies suggest that few mutations in TFs may be sufficient for changes in regulation (7, 16). Therefore, it is possible that even if the evolution of TFs is shaped by adaptation, we may found TFs to be less diverse than their TGs within a species. Additionally, lower diversity of TFs would be indicative of stronger selective constraints.

Recent advances in genome-scale sequencing and analysis, and readily available large-scale WGS datasets offer an exciting opportunity to investigate the evolution of regulatory networks over short time-scales *i.e.*, across strains or within species. Equipped with tens of thousands of sequencing runs on *E. coli* from various hosts and geographical regions, we set out to estimate the sequence diversity of a thousand genes. Using these datasets, we tested if transcription factors of a bacterial species are indeed more diverse than its target genes.

## Results

### I. Differences in nucleotide diversity can arise from differences in gene’s length

Previous studies have shown that TFs are less conserved - as measured by the presence/absence of orthologs - across bacteria (2, 9, 17). We hypothesized that this flexibility of TRN may be reflected in greater sequence variation of TFs over shorter time scales, *i.e.* across strains or within a species. As a corollary, and under the assumption that the difference in conservation across species between TFs and non-TFs is a reflection of positive selection, TFs may acquire more non-synonymous variation than their target genes (TG) within *E coli*. Also, under the assumption that TFs do not differ from their TGs in their underlying mutation rate and synonymous changes are under relatively weaker selection, TFs should be similar to TGs in their synonymous variation. Towards testing these ideas, we first performed a preliminary study of sequence variation in TFs and their TGs using a library of completely sequenced and assembled *E coli* reference genomes.

Starting from 614 reference genomes, we removed redundant genomes based on sequence similarity (see **Methods**) to assemble a final set of 123 genomes (Table S1, supplementary file 1). We obtained the experimentally verified TRN of *E coli* from the *RegulonDB* database (18). We excluded all interactions between TFs from this TRN to obtain a set of 142 TFs and their 1119 TGs (Table S2, supplementary file 1). We measured sequence variation of these genes across isolates in terms of nucleotide diversity (π), which is computed as the average pairwise nucleotide differences per base. Contrary to our expectation, we found no difference in nucleotide diversity of TFs and TGs (*P*_Wilcoxon rank sum_ = 0.054) (**Fig. 1A**). Then, we estimated nucleotide diversity separately from synonymous (π_*S*_) and non-synonymous (π_*N*_) sites. Again, both the estimates revealed no difference between TFs and TGs (*P*_Wilcoxon rank sum_ = 0.1065 for π_*S*_, 0.1099 for π_*N*_) (**Fig. 1B,C**). In the above analyses, we did not make use of the knowledge of regulatory interactions between these genes and only checked for the overall difference in TFs diversity when compared to that of TGs, TGs being representatives of average non-TF protein-coding genes. When we performed paired comparisons, where a TF was only compared with *its own* TGs, we found all of the above estimates of diversity to be lower for TFs. Surprisingly, even synonymous diversity of TFs was less than that of their TGs (*P*_Wilcoxon signed rank_ *=* 0.0008 for π, 0.0049 for π_*S*_, 1.6 × 10^−5^ for π_*N*_) (**Fig. 1D-F**).

**Fig. 1.**
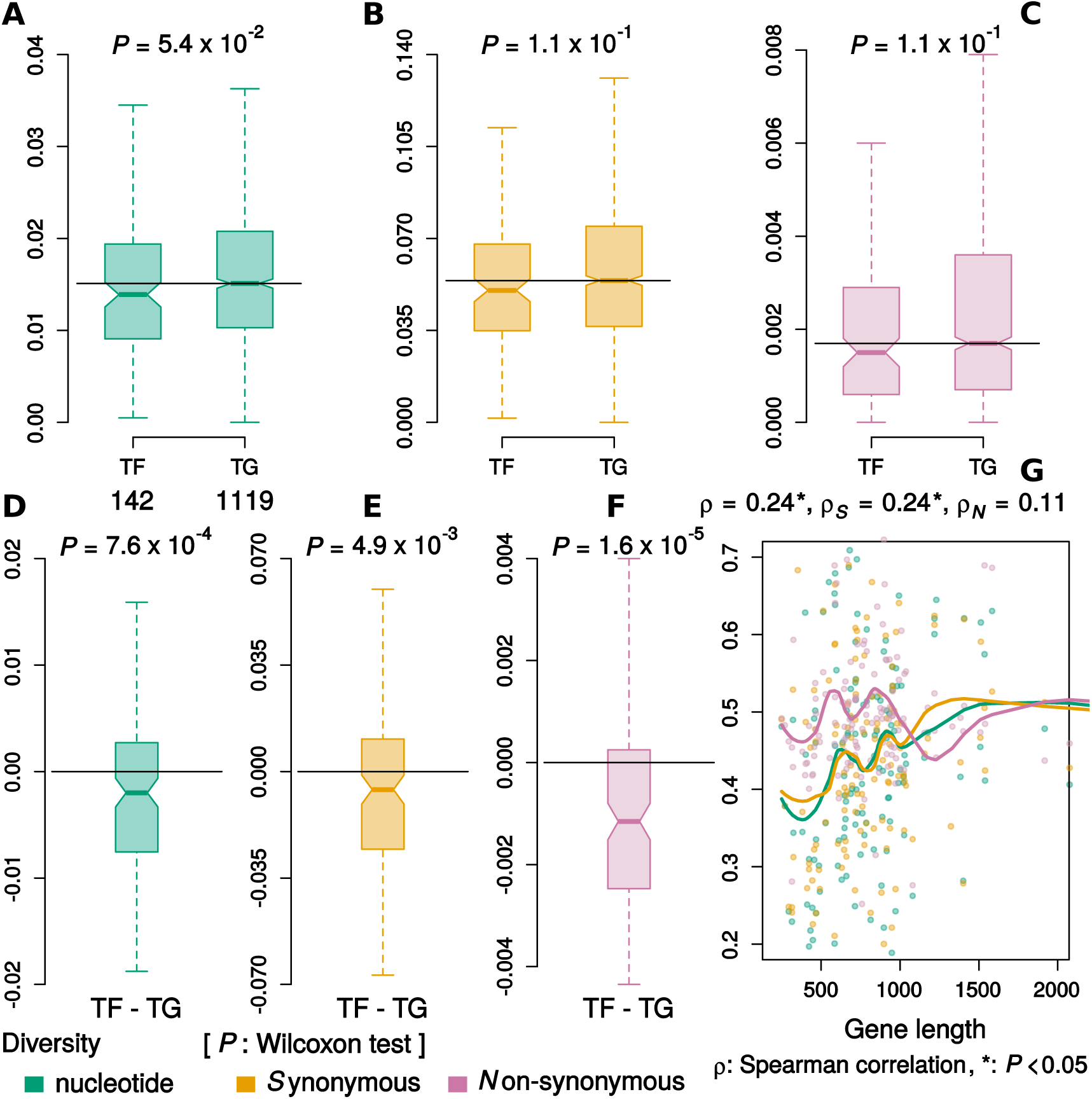
Sequence diversity and gene length.

**A-C**, Distributions of nucleotide, synonymous and non-synonymous diversity respectively, of TFs and TGs over 123 assembled genomes. **D-F**, Distributions of difference between the diversity of TFs and *their own* TGs for nucleotide, synonymous and non-synonymous diversity, respectively. Unpaired comparisons showed no significant difference but all paired comparisons did. **G**, Correlation between diversity difference (TF - TG) and TF’s length. Nucleotide (*P* = 0.0018) and Synonymous diversity (*P* = 0.0023) were positively correlated with gene length, whereas Non-synonymous diversity was not (*P* = 0.1016). Thick lines represent LOESS curves with a span of 0.5. Differences were Min-Max scaled for visualization. To improve resolution, outliers were excluded from box plots and y-axis for the scatter plot was restricted between 0.2 and 0.7 (which covers ∼93 % of data points).

Since the number of regulated TGs varies across TFs and ranges from a single TG to over 100 TGs, and we compared the mean of all TGs of a TF against its own diversity, TFs may show less variation than their TGs due to this large variation in sample sizes from which means were estimated. We addressed this issue by estimating the significance of our observation using a randomization test. Briefly, we simulated random networks by removing all TFs from the network and replacing original TFs with randomly selected TGs, keeping regulon sizes constant. In essence, this procedure removes all TFs, and instead considers a random set of non-TFs as TFs. We tested if non-synonymous diversity of TFs was lower than that of their TGs, in a paired comparison, for these simulated networks. We found that random networks rarely generated a difference as extreme as that observed in the original TRN (*P*_Randomization test_ = 0.001, 1000 trials).

TFs are generally shorter in length than non-TFs (Fig. S1, supplementary file 2). To test if this might be a factor in diversity estimate, despite the fact that the estimates are reported per base, we checked for correlation between these diversity estimates’ difference of a TF and its TG with TF’s length. Indeed, we observed a positive correlation for nucleotide diversity, as well as synonymous diversity. However, non-synonymous diversity did not show any correlation with gene length (*P*_Spearman_ *=* 0.0018 for π, 0.0024 for π_*S*_, 0.1016 for π_*N*_) (**Fig. 1G**). Consequently, when we excluded all regulatory interactions where TG was longer than the TF (leaving 107 TFs and 471 TGs), we found no difference between TFs and their TGs in nucleotide diversity and synonymous diversity, but a strong difference in non-synonymous diversity (*P*_wilcoxon signed rank_ *=* 0.2656 for π, 0.639 for π_*S*_, 2.62 × 10^−5^ for π_*N*_). Therefore, to test our hypothesis, we decided to estimate nucleotide diversity only in terms of non-synonymous diversity since gene length had no significant effect on the difference in non-synonymous variation of TFs and TGs.

### II. TFs are less diverse than their target genes within species

We expanded our analysis of nucleotide diversity across a limited set of 123 reference *E coli* genomes to a larger collection of short-read based genomes. These were sourced from publicly available sequencing projects covering clinical and environmental isolates of *E coli*. Since these datasets were larger than the earlier set and were expected to have greater redundancy, we used a different approach for filtering isolates (see **Methods**). We processed ∼16,000 sequencing runs from 24 projects and finally selected 15 datasets comprising a total of 2476 isolates (Table S3 and S4, supplementary file 1).

Nucleotide diversity is conventionally calculated from multiple sequence alignments of gene sequences extracted from assembled and annotated genomes. We developed a methodology that enabled us to use reads from WGS datasets to estimate nucleotide diversity. Briefly, we performed variant calling using *SPANDx* pipeline (19), inferred gene presence from the coverage, identified gaps and estimated diversity from SNP matrices generated by the variant calling pipeline. We also validated our approach and results using a small set of isolates for which both WGS data and assembled genomes were available (supplementary file 3). Since it is expected that the average nucleotide diversity would vary across samples, we standardized diversity estimates of all genes using the median and median absolute deviation (MAD) of diversity of TGs. This treatment would scale the median for TGs to zero and any difference in the diversity of TFs would appear as a shift in its median from zero, rendering diversity estimates comparable across samples.

We found that the non-synonymous diversity of TFs was less than that of *their own* TGs for all of the 15 datasets (*P*_Wilcoxon signed rank_ = 0.0018 – 3.4 × 10^−6^, after correcting for multiple testing) (**Fig. 2**). Therefore, we concluded that bacterial TFs acquire less non-synonymous variation than their TGs, irrespective of the variables like host, source, virulence and structure of a population.

**Fig. 2.**
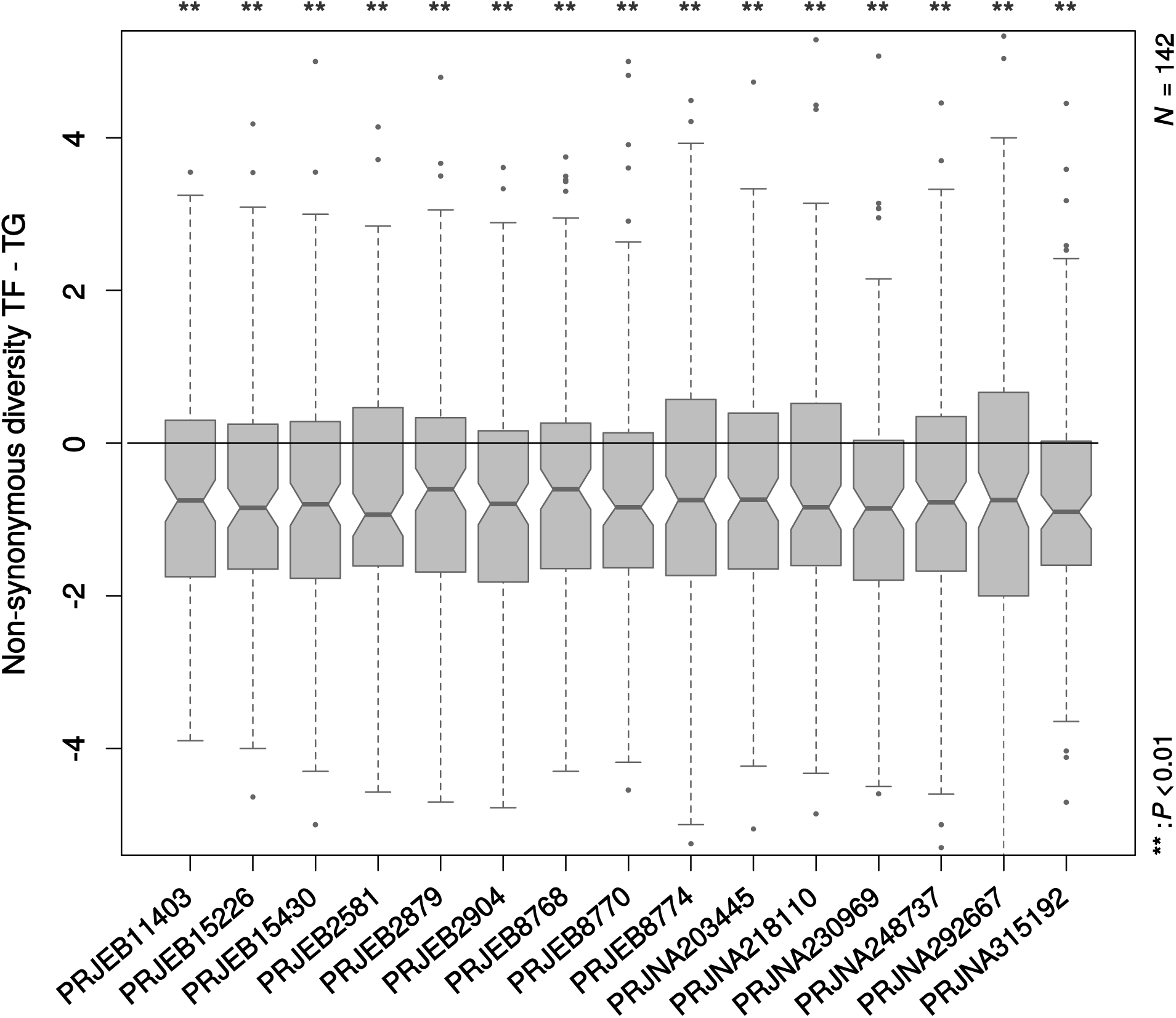
Non-synonymous diversity of TFs relative to their TGs.

For all 15 datasets, TFs showed less diversity than their own TGs, in a paired comparison. To make diverse datasets comparable, all values were rescaled using median & MAD of TGs, such that scaled median for TGs was zero. Y-axis represents the difference between scaled non-synonymous diversity of TFs and their TGs. P-values are based on Wilcoxon signed rank test of the hypothesis that TFs were less diverse than their TGs. To improve resolution, y-axis was restricted between −5 and 5 (which covers ∼ 96 % of data points).

### III. Genetic variation of TFs is constrained by their regulatory roles

The organization of the regulatory network is such that a few TFs regulate a majority of genes and are called global regulators (GR). In *E coli*, the 7 most prolific TFs are responsible for expression of 60% of the genes. These global regulators correspond to broad cellular programs such as carbohydrate and amino acid metabolism, respiration and growth, environmental sensing and stress responses (20). Since the targets of these global TFs belong to multiple functional categories, as defined in COG, these can also be called general TFs. Specific TFs, on the other hand, regulate target genes from a single pathway or at least from the same functional category (21).

The effect of mutations in a TF is expected to depend on its position within the TRN. TFs regulating a large number of TGs should be under stronger purifying selection. Consequently, we observed a negative correlation between TFs diversity (mean scaled non-synonymous) and its regulon size (as measured by the number of Transcription Units (TUs)) (**Fig. 3A**) (**ρ** = −0.5, *P*_Spearman_ = 1.64 × 10^−10^). The same observation was earlier made on variation across species (5, 22). Accordingly, the diversity of general TFs was significantly lower than that of specific TFs (**Fig. 3B**) (*P*_Wilcoxon rank sum_ = 1.15 × 10^−5^), tested with 8 general and 45 specific TFs in our dataset. However, even for these specific TFs, we verified that their diversity was lower than their TGs for all datasets (*P*_Wilcoxon signed rank_ = 0.032 – 0.001, after correcting for multiple testing) (Fig. S2, supplementary file 2). Therefore, we concluded that the diversity of a TF is constrained by the extent to which any change in TF can disturb the gene expression profile of the organism.

**Fig. 3.**
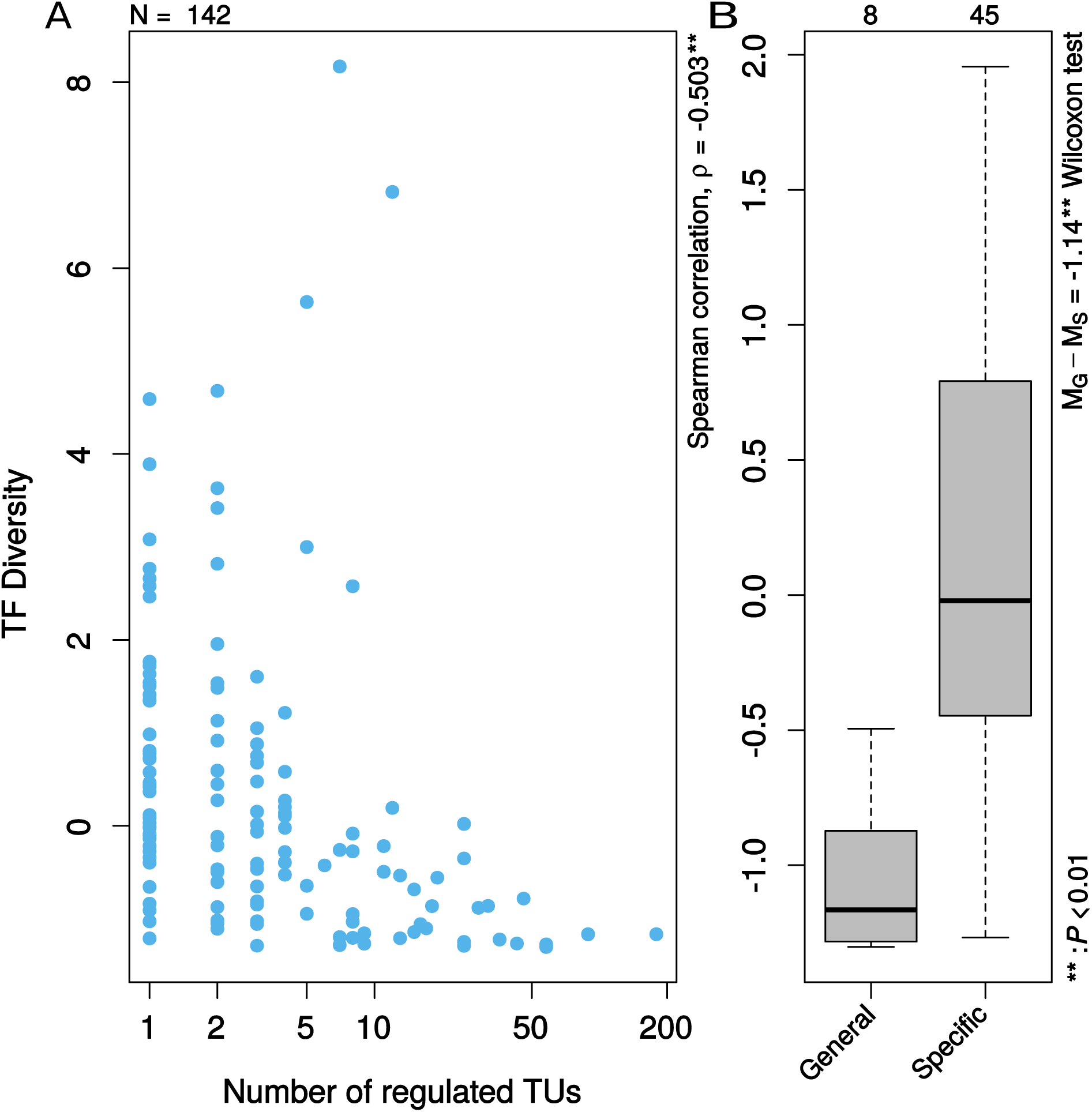
Regulatory constraints on diversity of TFs.

**A**, Non-synonymous diversity of TFs was negatively correlated with the number of regulated TUs (*P*_Spearman_ = 1.64 × 10^−10^). **B**, Accordingly, general TFs were less diverse than the specific TFs (*P*_Spearman_ = 1.15 × 10^−5^). To improve resolution, outliers were excluded from the plot. Y-axis represents mean scaled non-synonymous diversity of TFs. Scaling was done using median and MAD of TGs’ diversity for each of the 15 datasets and mean of the scaled diversity was taken over these datasets.

### IV. Conservation of TFs across species is also affected by their specific regulatory function

Previous studies had found TFs to be less conserved – in terms of presence/absence – than TGs (2, 17), across species and attributed these differences to duplication (4) and horizontal gene transfer (HGT) (5). However, the contribution of point mutations, if any, to the above observation remains largely unknown. In general, excessive polymorphism in a gene is indicative of weak selective constraints and thus, the gene is more likely to be eventually lost over long evolutionary distance. Under the assumption that these small, sequence-level, changes observed over short time-scales can explain gene’s presence/absence over longer time-scales, we had expected TFs to be more diverse in sequence than TGs within species. Contrary to our hypothesis, we found that the sequence diversity of TFs was lower than their TGs across isolates of *E coli*. We sought to reconcile these seemingly conflicting results by re-estimating conservation of *E coli*’s TFs & TGs across hundreds of bacterial species.

We performed bi-directional best hit based ortholog search using hidden Markov models of *E coli* TFs and their TGs across genomes of 246 species (Table S5, supplementary file 1) belonging to taxonomy class γ-proteobacteria which also includes *E coli* and *Salmonella*. We restricted our analysis to this class to increase our likelihood of finding orthologs for *E coli* genes. We defined “Conservation” of a gene as the fraction of species in which our search method could report an ortholog.

The underlying assumption of our hypothesis is intuitive and has not been formally tested in our knowledge. It is based on a postulate of the neutral theory of molecular evolution (23) that the same evolutionary processes govern variation within species and across species. Therefore, we first tested the validity of our hypothesis by estimating correlation between *Diversity within* species and *Conservation across* species, using all TFs and TGs. In accordance with our expectation, *conservation* was inversely correlated with *diversity* (**Fig. 4A**) (**ρ** = −0.42, *P*_Spearman_ < 2.2 × 10^−16^). However, it is evident from **Fig. 4A** that, for very low diversity, variance in *conservation* is much higher than that for the higher extreme. A possible explanation for this observation is that several genes, which are only present in a few bacteria, are most-relevant in their common natural habitat. *Conservation* of such genes across species is influenced more by HGT and duplication than by mutation accumulation. Therefore, in principle, TFs may be less *conserved* than their TGs across species despite being more conserved than their TGs within species. However, unlike previous reports, we did not find *conservation* of TFs to be significantly lower than that of their TGs across species (*P*_Wilcoxon signed rank_ = 0.116) (**Fig. 4B**).

**Fig. 4.**
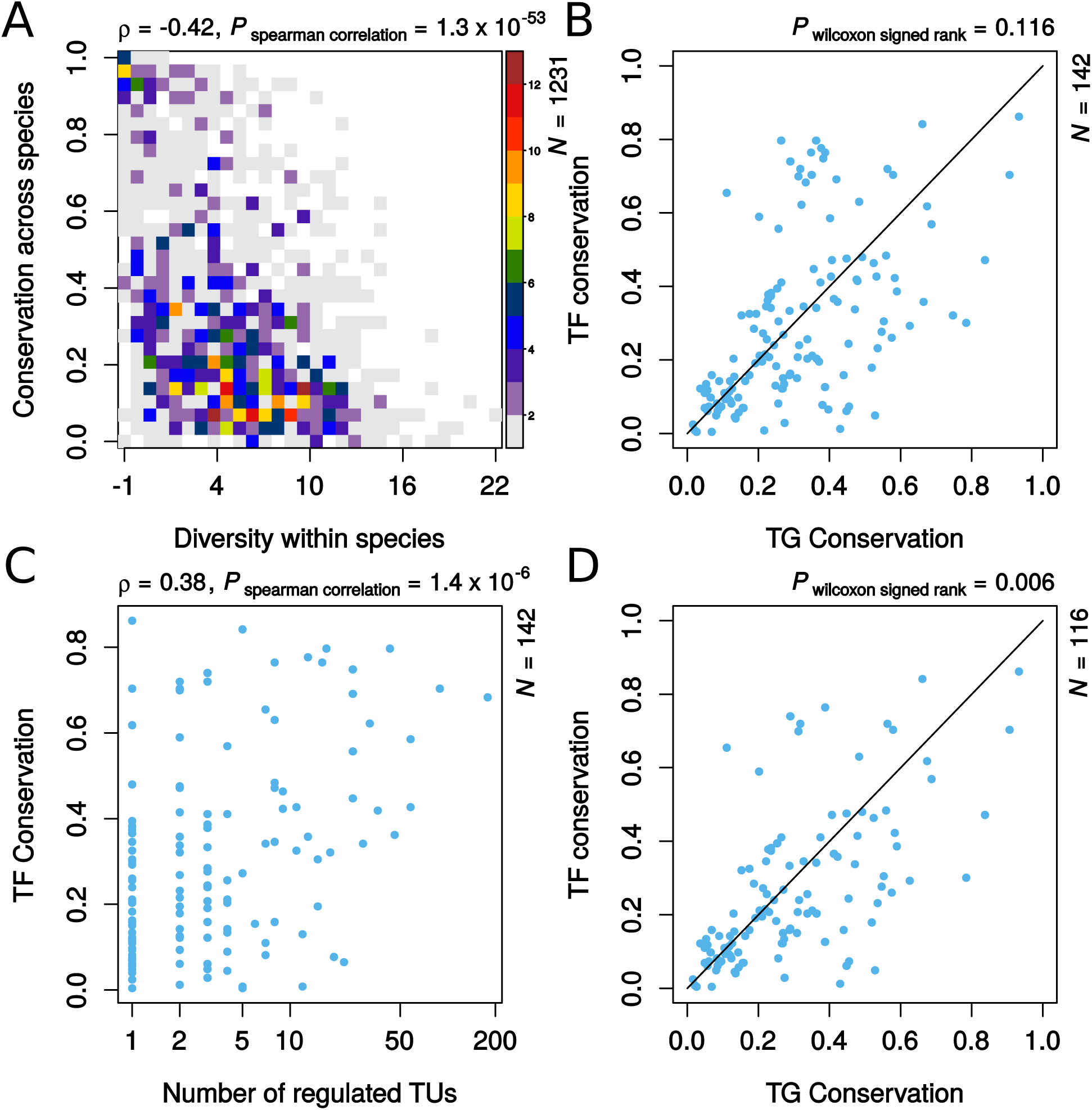
Effect of sequence diversity on conservation of TFs across species.

**A**, Conservation across species was negatively correlated with diversity within species. Y-axis represents fraction of 246 genomes with an ortholog for *E coli* TFs and TGs. X-axis represents scaled non-synonymous diversity averaged over 15 datasets. Scaling was done with median and MAD of diversity of TGs. Color scale represents count of data points in each bin. **B**, Conservation of *all of the* 142 TFs against that of their TGs. **C**, Conservation of TFs was positively correlated with the number of regulated TUs. **D**, Conservation of TFs, *excluding GRs*, was less than that of their TGs.

GRs were earlier found to be more conserved across species than other TFs (5, 22). Both of these studies were restricted to γ-proteobacteria. However, previous studies which reported low *conservation* of TFs estimated *conservation* across more distant groups of prokaryotes, and even eukaryotes, and did not find GRs to be highly conserved (2, 17, 24). Evolution of GRs was mostly vertical and least affected by HGT, unlike other TFs (5). This suggests that GRs might have evolved independently in distant lineages of prokaryotes (17). Since we restricted our analysis to the species of γ-proteobacteria, we also found *conservation* of GRs (*N*_TU_ >= 10) to be greater than that of other TFs (*N*_GR_ = 26, *N*_TF_ = 116 (Fig. S3, supplementary file 2) (*P*_Wilcoxon rank sum_ = 1.28 × 10^−4^). In fact, we observed a positive correlation between *conservation* of a TF and its regulon size (**ρ** = 0.38, *P*_Spearman_ = 1.4 × 10^−6^) (**Fig. 4C**). Consequently, excluding GRs, *E coli* TFs were less *conserved* than their TGs across species (*P*_Wilcoxon signed rank_ = 0.006) (**Fig. 4D**). However, these “local” regulators (LR) were also less *diverse* than their TGs within species, suggesting their relevance specifically to the natural habitat of bacteria.

These results, taken together, emphasize that even though TFs are generally thought to be less conserved than other genes, conservation of a TF seems to be affected by evolutionary distance between lineages, its position in TRN and its relevance to organism’s lifestyle (2), and a single statistical relationship is inadequate in capturing these complex interactions.

### V. TFs are enriched in mutations during adaptive lab evolution

In Lenski’s Long Term Evolution Experiment (LTEE) (25) and in several other adaptive lab evolution (ALE) studies, regulatory genes were reported to be mutated frequently (10). We had reported a higher frequency of mutations in regulatory genes in a prolonged stationary phase experiment compared to that observed in a mutation accumulation (MA) study (26). These results suggested that regulatory mutations contribute more towards adaptation than other mutations. Different runs of adaptation are expected to select for different mutations given the variability and complexity of natural environments. Therefore, we expected sequence diversity of TFs to be greater than that of other genes. However, as described above, we observed the opposite from our analysis using natural isolates. Since the above claim regarding regulatory mutations and adaptation was based on a few lab evolution studies, we decided to re-analyze these and other such studies using our methodology to test if TFs’ mutations during ALE are indeed more frequent than mutations in the other genes.

First, we identified 17 lab evolution projects with published (27–43) information about the experimental design, 4 of which were MA studies (40–43). For each study, we counted the number of non-synonymous mutations in TFs and TGs relative to the number of corresponding sites. 4 (29, 34, 35, 39) out of 13 ALE studies (27–39) had a significantly higher number of mutations in TFs than expected whereas none of the MA studies passed the significance test (**Fig. 5**) (*P*_One-sided Fisher’s exact_ < 0.05). Overall, ALE studies had greater odds of TF mutations’ enrichment than MA studies (*P*_wilcoxon rank sum_ = 0.024).

**Fig. 5.**
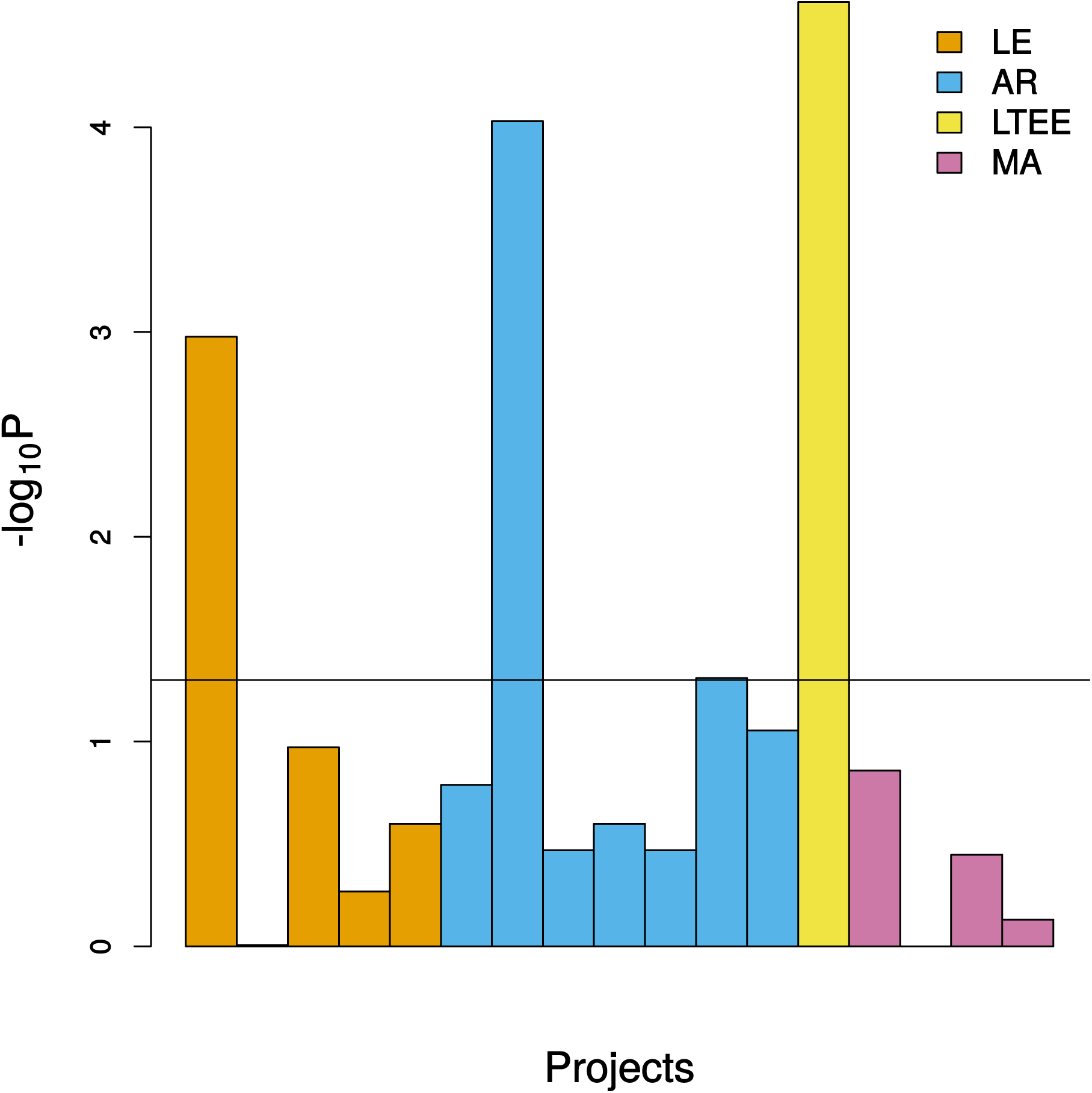
Relative frequency of mutations in TFs in lab evolution experiments.

Each bar represents a lab evolution study. Bars for adaptive lab evolution (LE) studies other than those selecting for antibiotic resistance are colored orange, the antibiotic resistance (AR) ones being shown in sky blue; Lenski’s LTEE is in yellow and reddish purple bars represent mutation accumulation (MA) studies. Height of bars represents transformed p-values of one-sided Fisher’s exact test of the hypothesis that the frequency of mutations in TFs relative to TGs is greater than expected from the ratio of sites. LTEE showed an enrichment for TF mutations. Majority of antibiotic resistance experiment didn’t show the same effect. Nevertheless, Odds Ratios for TF mutation enrichment were greater for ALE than for MA studies (*P*_Wilcoxon_ = 0.024).

Since our approach uses all of the observed mutation data and cannot consistently identify beneficial mutations across lab evolution studies due to differences in their experimental designs, we also tried a different approach based on literature survey. We identified those mutated proteins which the authors considered to have fitness benefits and counted the number of mutated TFs among 304 known and predicted TFs out of 4140 proteins in the reference genome. Here, we did not account for the site-count differences between TFs and TGs because the information on the actual number of mutations in a gene was not always available from the publication. In only 5 ALE studies (29, 33–35, 39), significantly more than expected (∼ 7%) mutated TFs were reported, 4 of which we had already identified (*P*_Fisher’s exact_ < 0.05) (Table S6, supplementary file 1).

2 of these studies (29, 39) had bacteria growing on minimal glucose, wherein the selection was for fast growth and presumably multiple adaptive paths were possible. 7 of the 12 ALE studies aimed to understand some aspects of antibiotic resistance (32–38), 3 of which (33–35) also showed an enrichment for TFs’ mutations. Depending on the mode of action of the targeted drug(s) in a study, mutations may only be selected in a specific enzyme, in which case, an excess of TF’s mutations would not be observed. This also holds for studies on auxotrophs (30). Often though, when a study focused on cross-resistance or on a drug with multiple mechanisms, selected mutations were more frequently among TFs (33, 34, 37) (Table S7, supplementary file 1). Other than the targeted enzyme, the resistance-conferring mutations often occurred in *marR, acrR, soxR* & *ompR*. These genes have an established role in conferring resistance, *via* a non-specific increased efflux activity, as a result of which, these are also common targets in selecting for cross-resistance (44). In (35), where bacteria were grown on a surface with regions of increasing concentration of a specific antibiotic, the selection might actually have been for fast growth since the chance of success dependent more on the timing of arrival to a higher concentration field than on the degree of resistance at that concentration.

In this section, we established that TF mutations are often enriched in ALE studies. The possible reasons for the lack of evidence for this effect in several other lab evolution studies are enzyme-specific selection pressure, small sample sizes and short duration. Lenski’s LTEE does not suffer from the above issues (25) and provides evidence for a high relative frequency of mutations in TFs over TGs. Another major issue with other studies analyzed here, in contrast to LTEE, was that they could only be used to test if TFs accumulated more mutations than TGs by the end of experiment. Due to their short duration and sequencing at limited time points, they could reveal nothing about if, and how, the relative strength of positive selection on TFs change over time. Therefore, we further explored mutation data from LTEE to derive an understanding of mutant frequency dynamics in natural populations.

### VI. Frequency of mutation accumulation in TFs declines over long-term evolution

A recent population-level study on LTEE (45) revealed the dynamics of molecular evolution over 60,000 generations with a 500-generation interval. This dataset enabled us to go beyond relative frequency estimation at a single time point and instead, observe the trajectory of these frequency changes over thousands of generations. It thus offered the possibility of reconciling our contradictory observations on TF vs. TG variation in experimental evolution and natural populations.

As reported earlier (46) and also in the above mentioned study, the rate of molecular evolution in LTEE is rapid and almost steady despite a decline in the rate of fitness gain. This rate was measured in terms of total derived allele frequency (DAF), “ *M* _*p*_ (*t*)=∑ *f* _*p*, *m*_ (*t*) for all mutations *m* in population *p* at time *t”* and *f* is the frequency of a particular mutation. Using this metric, we first tested if TFs accumulated more mutations than TGs. We averaged DAF of non-synonymous mutations for all TFs and TGs separately, assigning zero to non-mutated genes. For six non-mutator populations, we observed that DAF was significantly higher for TFs at all time points and a substantial increase was achieved in about first 10,000 generations (**Fig. 6A**).

**Fig. 6.**
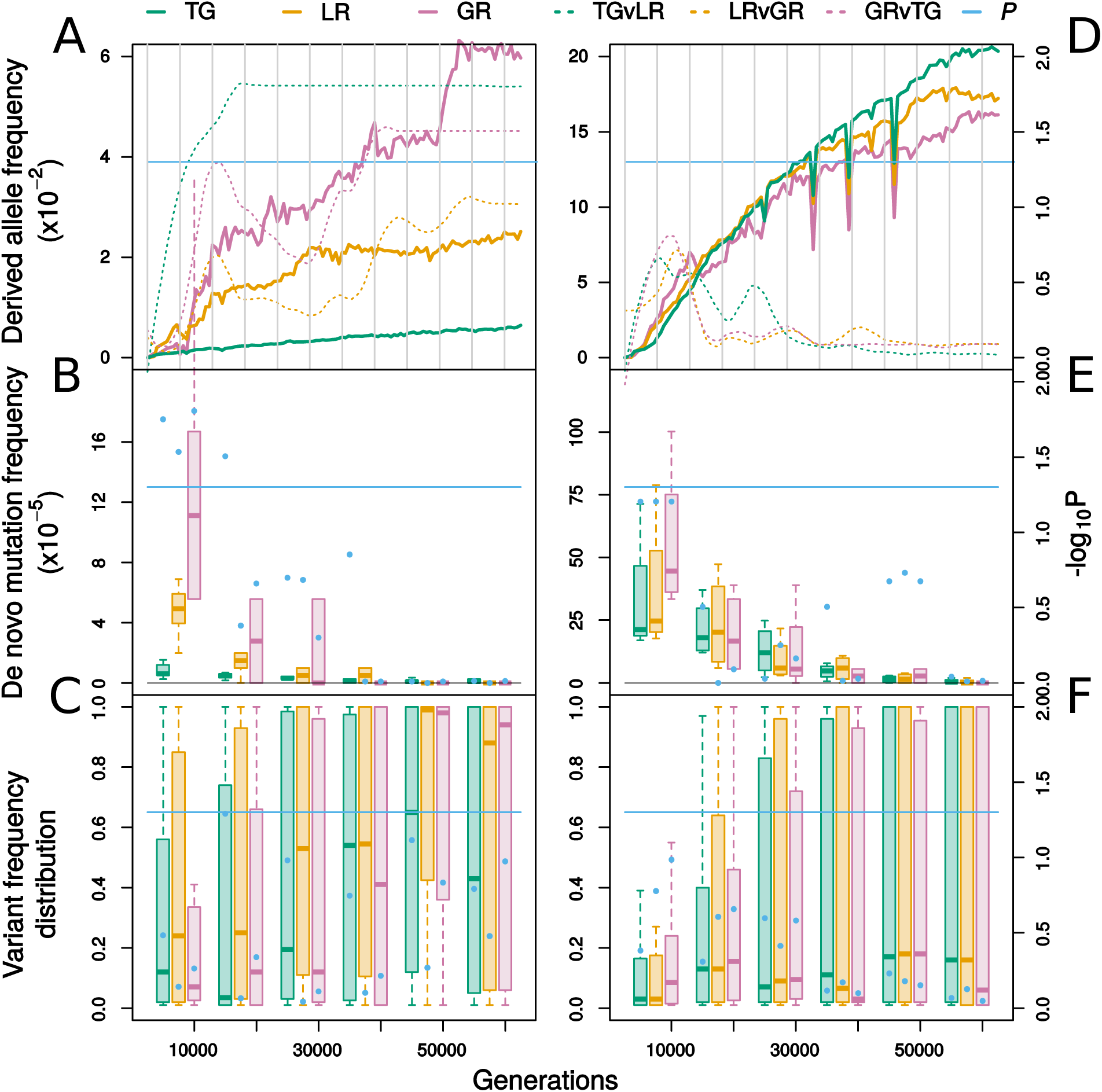
Distinct dynamics of molecular evolution for TFs and TGs in LTEE.

for six non-mutator populations: **A-C** and for four mutator populations: **D-F. A & D**, Average derived allele frequency of TFs and TGs over 60,000 generations. Thick curves represent averages over populations and broken curves show transformed p-values of three hypothesis: TG < LR, LR < GR, GR > TG. **B & E**, Distributions of count of *de novo* mutations appearing over intervals of 10,000 generations, normalized by number of sites. **C & F**, Distributions of maximum frequency attained by a variant within each interval. Variants were pooled from all populations. Blue dots show transformed p-values. Blue horizontal line marks the significance threshold (α = 0.05). Dots are in triplets, as are the boxes, such that they represent comparisons in the same order as above. P-values are based on Wilcoxon test.

The change in DAF can be brought about either by a change in the rate at which *de novo* mutations appear or by a change in the frequency these variants attain in the population. First, we plotted distributions of *de novo* mutations per site over 10K generation intervals for both classes (**Fig. 6B**). We chose this interval size since the smaller intervals didn’t capture mutations in all populations for all categories (TG, LR, GR). TFs accumulated more mutations than TGs only up to 20K generations. Then, over the same intervals, we pooled frequencies of all variants – existing and *de novo –* from all six non-mutator populations (**Fig. 6C**). Overall, TF variants didn’t reach significantly higher frequencies than TG variants. This may suggest that, on average, regulatory mutations were not any more beneficial than the mutations in target genes.

A majority of TF mutations in the first 10K generations were in fact in GRs instead of LRs (**Fig. 6B**). The significance of global regulatory changes in ALE has been noted previously (10, 11, 47). Global scale changes in gene expression pattern are required for a cell to achieve an optimal metabolic flux state, on which its relative fitness depends, since transcription is costly due to limited availability of RNA polymerase (48). Even single mutations in the hubs of regulatory networks can achieve this goal, by simultaneously increasing the expression of genes required for success in the testing environment and “switching off” the parts of network irrelevant in the present context. What is interesting to note here is that the frequency of *de novo* mutations in GR showed a much faster decay than that for other genes. Moreover, the early stage GR mutations did not attain high frequencies within the first 10K generations, presumably due to clonal interference (**Fig. 6C**). However, at least a few of these mutations reached fixation by the end of 60K generations.

The above analysis was on mutations per site basis, without taking into account the differences in mutation propensity of individual genes. In the extreme case where all of the GR mutations targeted a single gene, the observed trend cannot be generalized. Therefore, we performed the above analysis after removing the extreme outliers gene (one which had many mutations with a large fraction reaching fixation) from each category (TG: *pykF* = 7/10, LR: *iclR* = 5/8, GR: *arcA* = 4/6) (**Fig. 7**). TG mutations other than in *pykF* only reached low frequencies in the first 10K period, even slightly lower than those in GR. Excluding *arcA*, GR were not different from LR in their mutation frequency. Moreover, most of the GR mutations reaching fixation were in *arcA*. The frequency distribution of LR was least affected and was in fact, higher than that of GR for the greater part of 60K generations.

**Fig. 7.**
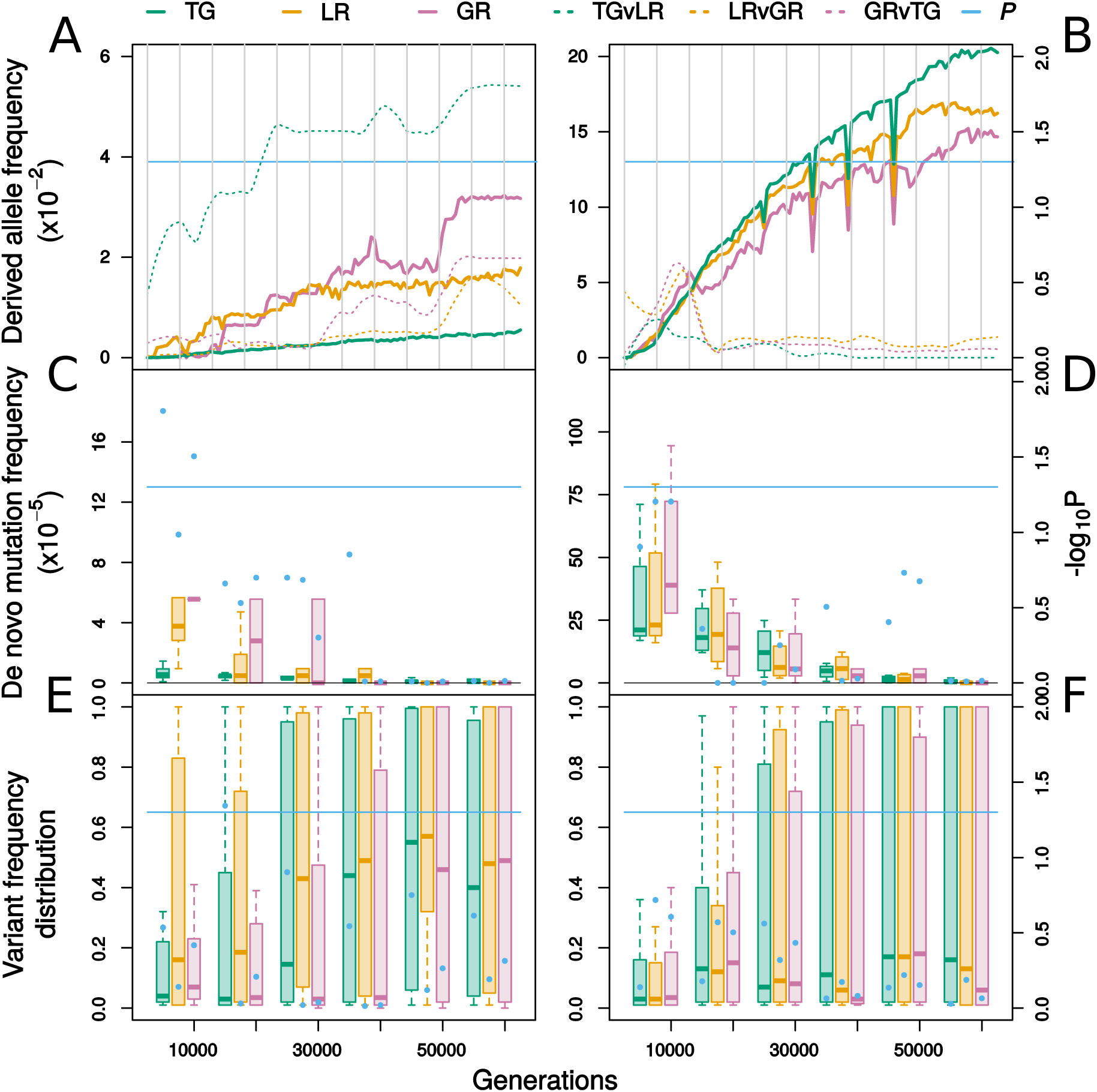
Differential dynamics of molecular evolution for TF and TG in LTEE, same as in **Figure 6**, except that an extreme outlier gene (one with many mutations and a large fraction of mutations reaching fixation) was removed from each class of proteins.

Since we didn’t have information on the linkage of variants, we could not use this data to directly estimate nucleotide diversity at non-synonymous sites. However, estimates of *de novo* mutations frequency and the distribution of population frequency of the variants, when taken together, offers a qualitative approximation of the relative diversity of the two classes. In the first 10K generations, we expect TF diversity to be greater due to high frequency of *de novo* mutations. From 10-20K generations, diversity of TGs was likely to be lower than that of TFs, since many of these variants were present at frequencies close to zero. By the end of 60K generations, population frequencies of variants of both classes were similar but only TGs were still accumulating *de novo* mutations. At this point and beyond, TG diversity may be equal to or even higher than the TF diversity.

Above results were based on six non-mutator populations (Ara-5, Ara-6, Ara+1, Ara+2, Ara+4, Ara+5). We did not find these results to hold for four mutator populations (Ara-2, Ara-4, Ara+3, Ara+6) (**Fig. 6, D-F**). In contrast to the non-mutator populations, a majority of mutations in mutator populations are non-beneficial. In such a regime, the patterns of mutation accumulation are expected to be governed by the differences in selective constraints on various genes. By the end of 60K generations, TGs in mutator populations appeared to have accumulated as many mutations, if not more, than in TFs (**Fig. 6D**), unlike in non-mutator populations. Besides, towards the end, TF and TG variants in mutator populations differed less in their frequencies than in non-mutator populations (**Fig. 6F**). A majority of variants in both classes were likely neutral to slightly deleterious such that their trajectories were governed by genetic drift.

To further test that the high frequency of TF mutations in non-mutator populations were due to positive selection, we analyzed one of the very few MA studies to have sequenced isolates at multiple time points (49). ∼36 clones were sequenced at 6 time points spanning ∼8,000 generations. At all time points, the fraction of sites mutated was higher for TGs than TFs, contrary to our observation in LTEE (Fig. S4, supplementary file 2).

Unlike other lab evolution experiments which only offered a snapshot of molecular evolution processes, LTEE provided us a record of the dynamics of variation within TFs and TGs spanning 60,000 generations. Even so, nature’s evolution experiment has traversed millions of generations and has likely passed through multiple fitness peaks unlike LTEE, which is yet to reach one. It is remarkable then, that even in the relatively short duration of this lab experiment, we could already observe TFs mutational frequency falling to the level of that of TGs. Especially in the mutator populations, where most mutations were likely to be neutral to slightly deleterious, this suggests a relatively faster deceleration of mutation accumulation in TFs over TGs. Besides, variants did not significantly differ in the frequencies they reached in the population. We extrapolate from these trends that TFs would acquire less mutations than TGs, as the experiment continues, and in principle, after millions of years, one would find TFs to be less diverse than their TGs.

## Discussion

We showed that bacterial TFs are less diverse in sequence than their TGs within species, *i.e.*, across short time-scales, and that their diversity is a function of their regulon size. It has been reported previously that global regulators (GR) are more conserved across species than other TFs (5, 22). However, even after excluding GRs, we found that TFs were more conserved - in sequence - than their TGs within species. This was contrary to the conservation of these “local” regulators (LR) – in terms of presence/absence – across species.

If two bacterial species have widely different environments, then their set of TFs are also expected to be different. However, within species, the niche differences may not be drastic enough to warrant diverse TF alleles. Under this scenario, the low sequence diversity of TFs within species is indicative of stronger selective constraints, imposed by the requirement of their optimal activity in a given environment. As a corollary, adaptation to a new environment may demand a new optimum of gene expression which is conferred through mutations in TFs. Indeed, multiple adaptive lab evolution (ALE) studies were found to be enriched with regulatory mutations (10). We performed a statistical analysis on many experimental evolution studies to verify this observation. Indeed, we found that TF mutations were enriched in ALEs under those selection pressures which can be satisfied by changes in multiple pathways. In contrast, none of the mutation accumulation (MA) experiments showed an excess of mutations in TFs.

To observe the long-term dynamics of the above trend, we analyzed whole-population data from an evolution experiment spanning 60,000 generations (45). We found that the frequency of mutations in TFs rapidly rose above that of TGs in first 10,000 generations and then declined over time. This trend was stronger for GRs and the decline was faster. However, only mutations in a few specific GRs conferred an advantage whereas multiple LRs were found to have beneficial mutations. In mutator populations, TFs and TGs accumulated mutations at similar rates and towards the end, any difference in trends seemed to be in accordance with selective constraints.

Based on our observations on diversity of TFs across time-scales, and the existing body of literature, we put forward the following model of TRN evolution in prokaryotes (**Fig. 8**). As a population first encounters an environment, it experiences global expression changes, brought about by mutations to regulatory hubs (11). Since these changes may also have adverse pleiotropic effects, as evolution proceeds, more mutations accumulate in LRs (47). As a consequence of these mutations, specific pathways of TRN, which are irrelevant to the present selection pressure, are inactivated (50). However, the fitness benefit of these TF mutations decline over time, likely as a consequence of diminishing returns epistasis (51), and as adaptation decelerates, selective constraints play a bigger role in the mutation frequency. In a well adapted population, the optimal variants of TFs are maintained by purifying selection. Across environments, different segments of the TRN are targeted and inactivated, such that over millions of years, species adapted to different environments have few TFs in common (2).

**Fig. 8.**
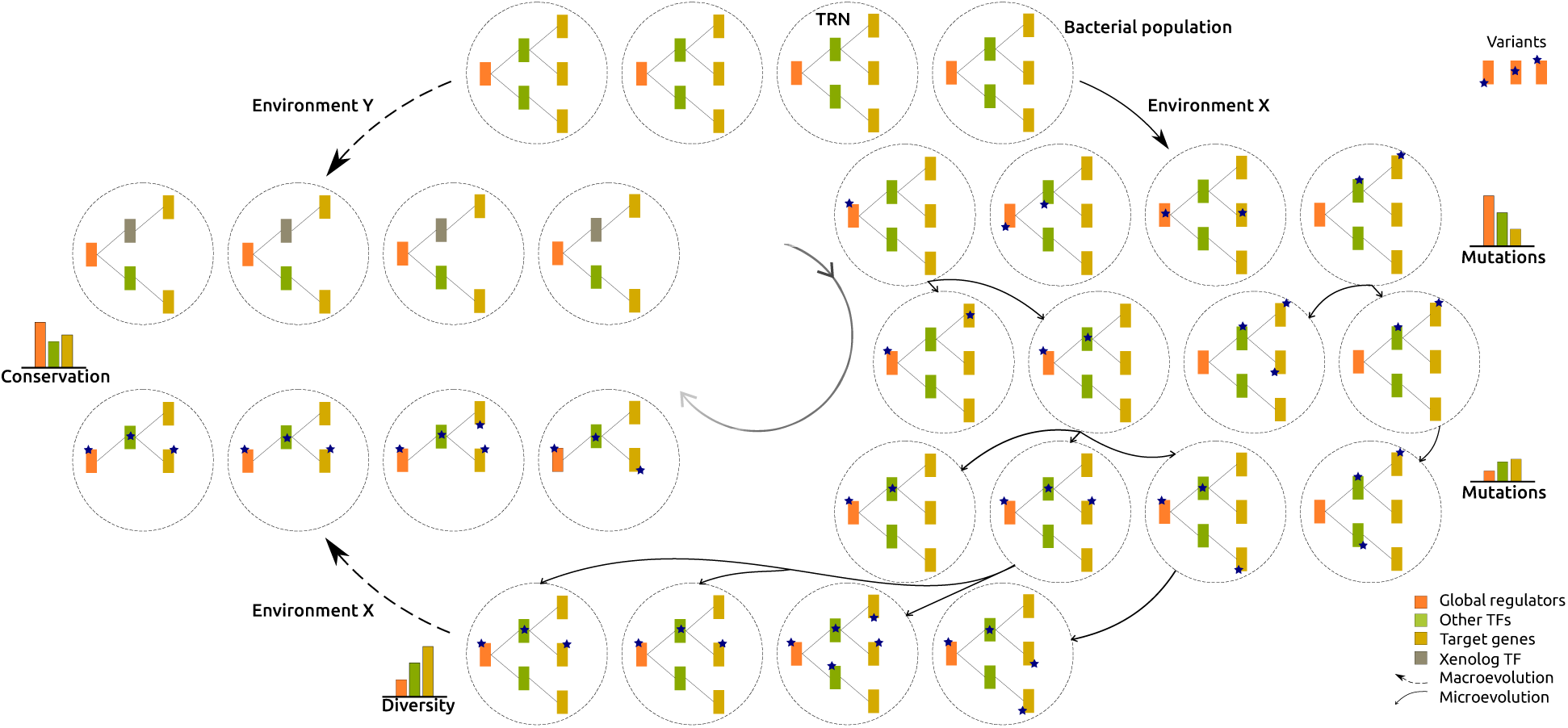
A graphical representation of the proposed model of TRN evolution in Bacteria.

A population exposed to an environment X rapidly accumulates mutations in TFs, especially GRs. Some of these mutants reach fixation while also accumulating mutations in other TFs and TGs. In long-term, mutations in local TFs are more beneficial than in global TFs. Farther out in time, when the population is well adapted, mutants of greater fitness rarely appear and hence, TFs show low sequence diversity. Also, irrelevant TRN modules are eventually lost. In a different environment Y, adaptation may proceed through integration of or substitution by xenologs in the native network. Thus, across species comparisons show low conservation of TFs relative to their TGs.

The above model does not underestimate the significance of HGT and duplication in the growth of regulatory networks. However, it emphasizes the role of small-scale changes, observed over short time-scales, in its modification. Specifically, in the early stages of adaptation, these changes set the path for long-term evolution of the network and facilitates pruning of branches irrelevant to the new environment. A major concern with this proposition might be extendability of results observed in LTEE, which is unrealistically simple as opposed to natural environments. Rampant HGT in nature is likely to have a strong effect on diversity of genes. Besides, the known range of selection pressures which lead to TF mutation enrichment is limited due to a prevailing bias towards studying evolution to antibiotic resistance. Large-scale evolution experiments, with more complex environments and employing diverse selection pressures, would be the true tests of importance and dynamics of mutations in TFs in the course of adaptation.

## Materials and Methods

### I. Data acquisition

The meta-data table for *E coli* WGS reads datasets, generated on Illumina platforms, was acquired from NCBI SRA database (http://trace.ncbi.nlm.nih.gov/Traces/sra/sra.cgi Accessed July 2017). This table included 49680 sequencing runs from 2114 bioprojects. The projects were analyzed in the descending order of the number of runs. For any given project, the meta-data table was downloaded from EMBL-EBI European Nucleotide Archive (https://www.ebi.ac.uk/ena). The runs were selected based on the following criteria:-library layout: “Paired”, library source: “Genomics”, library selection: “Random/ unspecified/ size fractionation/ random PCR/ PCR”, Coverage:- “>= 50X” and were downloaded from their respective FTP addresses mentioned in the ENA meta-data table using aria2 (https://aria2.github.io/). Variant calling was performed on 24 projects with 16116 clinical and environmental isolates in total. 15 of these projects, which had at least 50 sequencing runs left after declustering [see IV], were selected for further analysis.

The information on regulatory interactions among *E coli* proteins was acquired from the RegulonDB database (v9.4) (18). A set of 146 experimentally verified TFs and 1,119 TGs regulated by these TFs was extracted for the analysis, after excluding those genes for which multiple sequence alignment, based on the selected Refseq genomes (52) [see **V**], showed gaps in the sequence corresponding to the reference genome (*E coli* K-12 MG1655), arguing that these gaps can represent additional domains which may result in a different protein function. All regulatory interactions among TFs were excluded from the network which reduced the set of TFs to 142.

### II. Variant Calling

SPANDx v3.2 pipeline (19) was used for variant calling with default settings. The original script was slightly modified for ease of integration into our custom pipeline. *E coli* K-12 MG1655 genome (NCBI nucleotide database accession = NC_000913.3) was used as a reference for read alignment. The output SNP matrix was processed to remove low quality variants, variants for which base call was ambiguous in > 10% runs, and those outside protein-coding regions. Remaining ambiguous bases were replaced with the most-frequent base at the position, following which if the position had no variation, than it was excluded.

### III. Coverage-based gene detection

Presence of genes were detected separately from the variant calling pipeline based on the breadth and depth of sequencing coverage. “Breadth” implies the fraction of gene length which was covered by at least one sequencing read and “Depth” implies the average number of reads that mapped to each position of the gene. These quantities were estimated using *bedtools coverage* (v2.25.0) (53) and *samtools bedcov* (v1.3.1) (54) (with mapping quality > 50) respectively. First, genes above a minimum breadth of 0.6 were selected. Then, per base depth was calculated by dividing the total read base count by gene length. Genes with zero average base depth were excluded. A mean over all remaining genes, 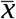, was calculated and 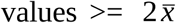 were excluded to obtain a Gaussian-like distribution of depth coverage with mean, 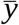, and standard deviation, *s*. Genes with depth 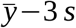 were considered present in the isolate. The runs with < 3000 genes were excluded. Only the variants corresponding to the detected genes were retained.

### IV. Selection of runs from WGS datasets

Sequencing projects, especially the ones with thousands of runs, were expected to contain many highly similar isolates. The redundancy in these datasets could bias our results. Therefore, a declustering step was employed. From the processed SNP matrix, lists of codon variants were generated for ea ch run. Codon distances (fraction of variant codons) from the reference genome were calculated for 125 genes which were present in all of the 15840 analyzed sequencing runs. These genes were selected from a set of 1710 core genes of 123 Refseq strains [see **V**], excluding genes with gaps in multiple sequence alignment of these strains, keeping only the genes with nucleotide diversity (based on 123 strains) > 75th percentile and <= IQR + 75th percentile. Codon distances were used to map a set of runs onto a 125-dimensional euclidean space, the corresponding distance matrix was generated and a subset was selected such that the minimum distance between any two runs was greater than 0.1.

### V. Selection of assembled genomes

Completed genome assemblies were downloaded for 614 *E coli* genomes from NCBI Refseq database (Accessed Oct, 2018). Since there was redundancy in this dataset as well, strains were selected using Mark Achtman’s MLST scheme as follows. The 7 gene fragments (*mdh, gyrB, recA, icd, purA, fumC, adk*) corresponding to *E coli* K-12 MG1655 were used in BLASTn (55) as queries against the above target genomes and the best hits with max *E* = 10^−5^, at least 70 % identity and 90 % overlap were identified. Strains with any fragment missing were removed. A sequence of whole number percentages of identity and overlap was created for each strain and only 1 randomly selected representative of each sequence was retained. This procedure ensured that the 123 selected strains differed by at least 1 % from others in their identity and overlap on at least 1 of the 7 gene sequences.

### VI. Nucleotide diversity estimation from WGS reads

Using the lists of codon variants from selected runs, “pseudo-codon” alignments were generated by initializing each row of the alignment with the reference gene sequence and then substituting reference codons with variant codons. Only those runs where the gene was detected were included in the alignment and the runs with an intermediate stop codon or with any mutation at the reference stop position to a non-stop codon were removed. Any variant codon with missing site(s) was considered entirely missing and all columns with gaps were removed from the alignment. From these alignments, nucleotide diversity was estimated conventionally, as the average pairwise nucleotide difference per unit length of the gene.

### VII. Nucleotide diversity estimation from assembled sequences

Protein orthologs were identified across strains with a custom-script using BLASTp (v2.2.29+) with minimum percent identity and query coverage of 50%, *E* < 1×10^−5^ and BLOSUM80 as the substitution matrix. Bi-directional Best hit criterion (56) was applied and ortho-groups were classified as strict core - members of which were found in all of the 123 genomes, closed group - every member of which could identify all other members, and open group - some members were not identified by other members. In case of open groups, only the “closed’ part was retained. Corresponding protein and nucleotide sequences were extracted and multiple sequence alignments of protein sequences were generated using CLUSTAL OMEGA (v1.2.1) (57). These protein alignments were converted to codon alignments using PAL2NAL (v14) (58). All gapped-columns were removed and nucleotide diversity was estimated in the conventional way as the average pairwise difference per site.

### VIII. Regulon diversity estimation

For a paired comparison of TF and TG diversity, the diversity of the regulon corresponding to the TF was estimated in the following manner. First, mean diversity of each transcription unit (TU) regulated by the TF was calculated. One or more genes when transcribed together on a single mRNA under a single promoter represent a transcription unit (TU). A single operon may have multiple fully or partially overlapping TUs. For the fully overlapping TUs, only the largest one was considered. Since the length of operon (and TU) is a function of the length of the corresponding metabolic pathway and we wanted to eliminate the effect of pathway lengths on diversity estimates, the grand mean diversity of TUs, instead of the weighted mean, was calculated for each TF.

### IX. Ortholog detection across species

For the analysis of conservation of proteins across species, 246 Uniprot reference proteomes (Accessed in Sep. 2016) from class γ-proteobacteria were used. Phmmer (HMMER 3.1b2) (http://hmmer.org/) with max *E* = 10^−6^ along-with Bidirectional Best Hit criterion was used to identify orthologs of reference’s TFs and TGs across these proteomes. Pairwise global alignment of hits with reference proteins was done using needle (EMBOSS:6.5.7.0) (59) and hits with > 10% alignment gaps were rejected. For a typical TF which is about 250 amino acids long and has a DNA-binding domain (DBD) of about 20 residues, this threshold improved the odds that it had the same DBD as the reference and likely performed an analogous function.

### X. Statistical Analysis

Distributions of Diversity and Conservation for TFs were compared against corresponding distributions for TGs using one-sided Wilcoxon rank sum test and Wilcoxon signed rank test for unpaired and paired comparisons respectively. Non-parametric tests were used because the underlying distributions were unknown and skewed. Correlations were tested using Spearman correlation test since a linear relationship was not assumed *a priori* and the interest was in assessing if the relationships were monotonously positive or negative. Mutation enrichment in TFs in lab-evolution projects was tested using one-sided Fisher’s exact test. P-values were corrected for multiple testing with Holm-Bonferroni correction. All statistical tests were performed using the statistical programming language *R* (v3.4.4).

## Availability

In-house scripts for analyses done in this study along with required input files are available at GitHub (https://github.com/A-Farhan/sequence_diversity_ecoli_TFs).

## Supplementary Data

1. File 1 - (xls) - Supplementary tables
2. File 2 - (pdf) - Supplementary figures
3. File 3 - (pdf) - Validation of nucleotide diversity estimation using WGS reads

## Acknowledgments

We thank Deepa Agashe, Sunil Laxman and Sabarinathan Radhakrishnan for reading and offering very helpful comments on an earlier version of this manuscript. We are also grateful to Deepa Agashe for mutation accumulation time-series data. We’d also like to show our gratitude to Sankarshan Talluri and the late Pragati Dembla for valuable discussions throughout the course of this work.

