## supplementary file 2 for "Dynamics of genetic variation in Transcription Factors and its implications for the evolution of regulatory networks in Bacteria"

**Supplementary Figures**

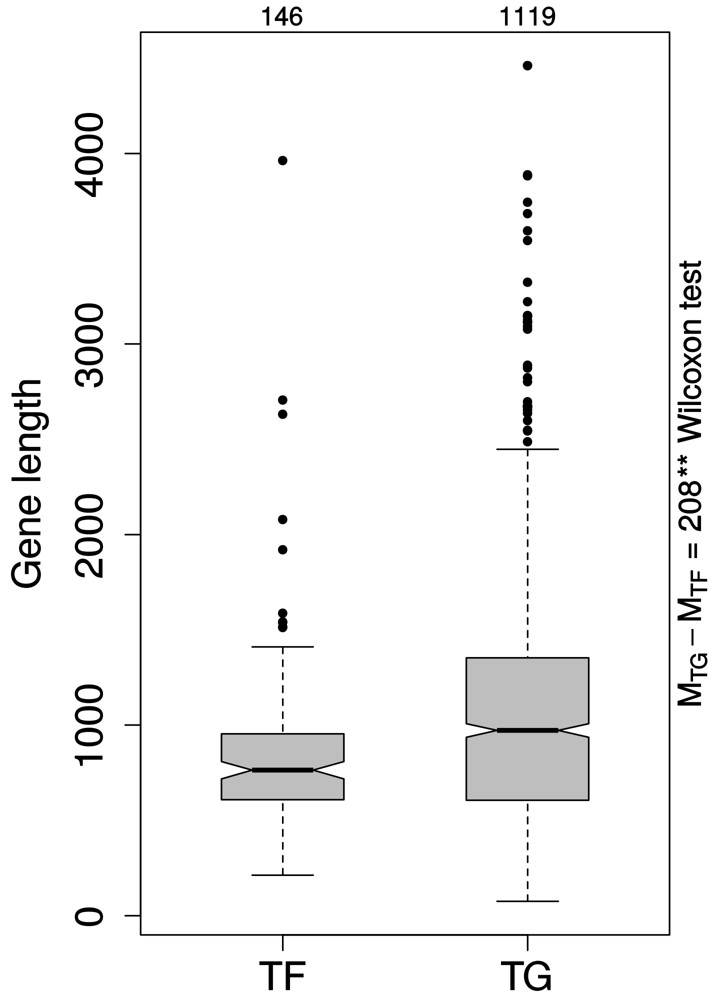


Figure S1. Gene length difference between TFs and TGs. TFs are significantly smaller than TGs ($P_{\text{Wilcoxon rank sum}}$= 9.31 x $10^{-6}$).

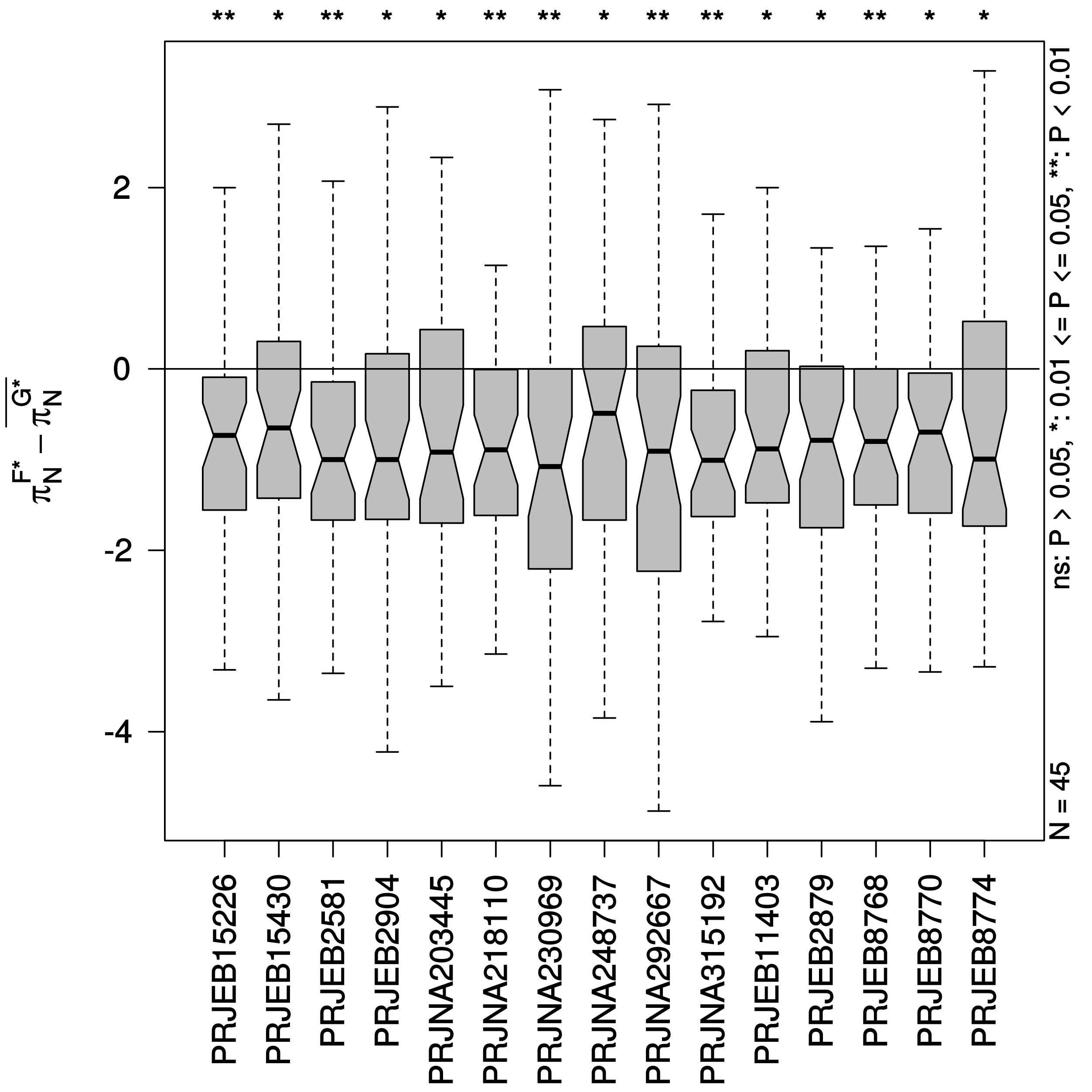


Figure S2. Non-synonymous diversity of specific TFs and their TGs. Distributions of differences in the scaled non-synonymous diversity of specific TFs and their TGs. As for the entire set of TFs, specific TFs were also less diverse than their TGs, for all 15 datasets, despite regulating single pathways or at best one functional category. P-values are based on Wilcoxon signed rank test.

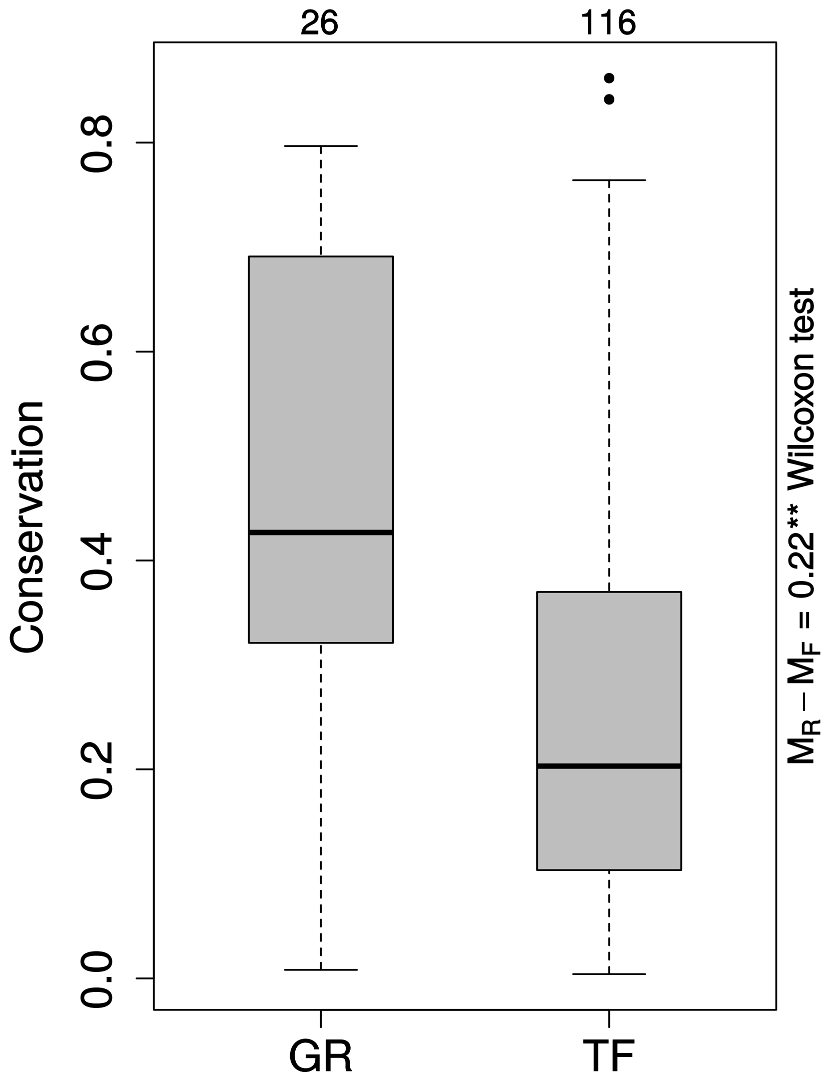


Figure S3. Conservation of GRs and other TFs. Conservation distribution of GRs in comparison with other TFs. Clearly, GRs are more conserved within γ-proteobacteria ($P_{\text{wilcoxon rank sum}} = 1.28 \text{ x } 10^{-4}$).

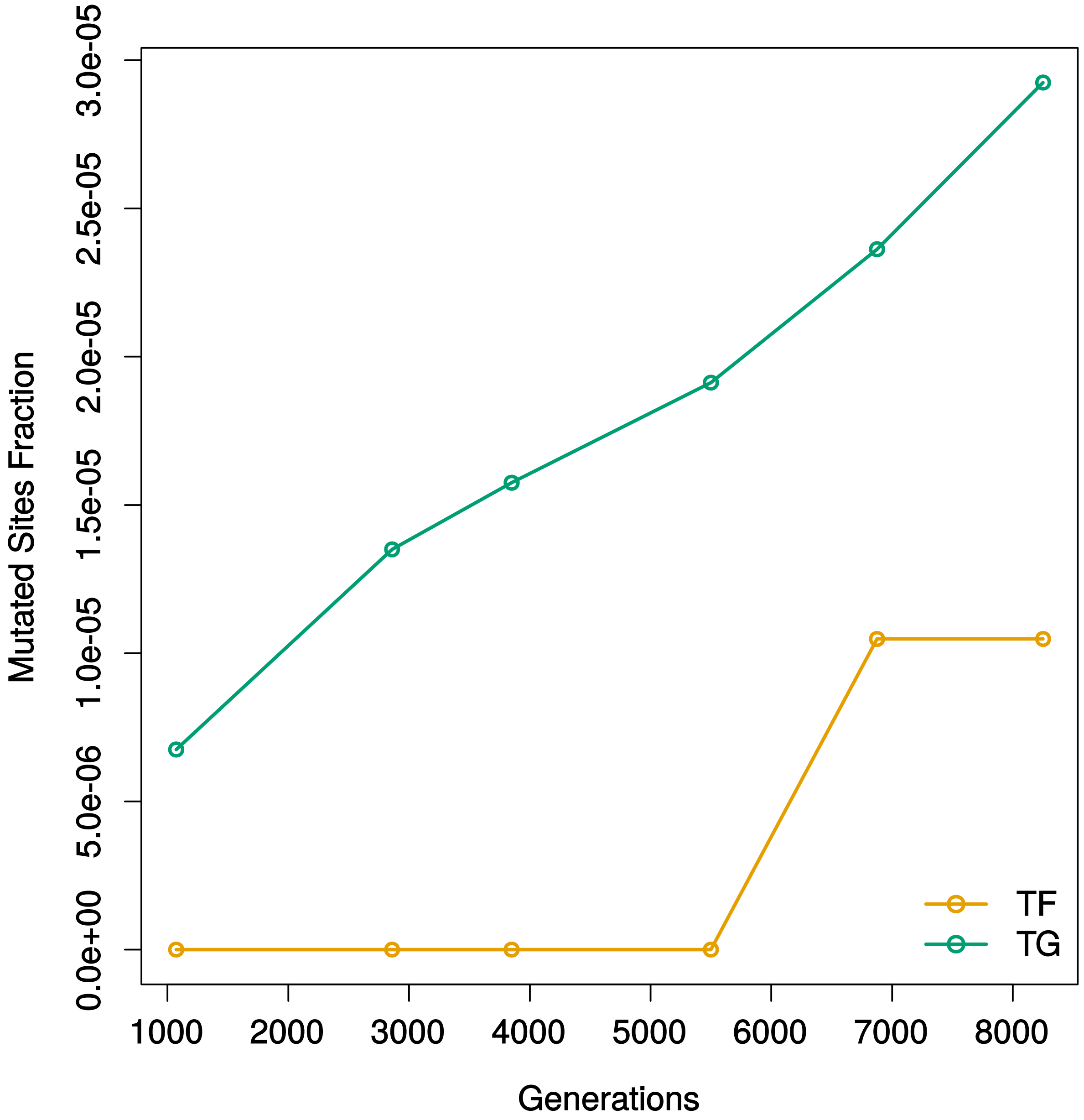


Figure S4. Relative frequency of mutations in TFs and TGs in MA. Change in frequency of observed mutations in TFs and TGs over 8000 generations of mutation accumulation. In stark contrast with LTEE where mutation frequency in TFs rapidly increased above the level of TGs during first 10,000 generations, TFs did not attain, at any point, as high frequencies as those for TGs in MA.
