## supplementary file 3 for "Dynamics of genetic variation in Transcription Factors and its implications for the evolution of regulatory networks in Bacteria"

### Validation of Nucleotide Diversity estimation using WGS reads

We used a read-mapping and variant calling approach to detect variation in the coding sequences of *E coli* genes across divergent strains. More conventionally, the sequence diversities are estimated from Multiple Sequence Alignment of orthologs. The conventional approach would limit us to assembled and annotated genomes. Out of 614 genomes available on NCBI Refseq database, we were left with only 123 strains for our analyses, after removing the redundant ones. In contrast, our approach, which is also scalable, allowed us to expand our search to a much larger number of strains available in public database for which genome assembly is not available. Whether the 123 strains make a fair representation of the *E coli* population and if the sample size is sufficient to detect our effects of interest is a problem that we have not specifically addressed here but our objectives require us to be able to quantify sequence variation across several datasets, including but not limited to those of experimental evolution, for which assembled genomes are neither available nor required.

We expect to reach to the same conclusions using our approach, as the conventional approach on the assembled genomes. To validate our results, we identified 24 strains for which WGS raw data was also available. We performed our analyses on these strains using the mapping approach as well as the MSA approach. We tested the strengths of correlations of various measures between these two methods, referred to as the “reads approach” and the “assemblies approach” to assess the accuracy of our method to estimate sequence diversity.

We obtained a strong correlation between the reads approach and the assembly approach for gene detection, albeit with a slight overestimation of gene count (Figure V1) ( $R^2$ = 0.91, slope = 0.93, simple linear regression). As for the nucleotide diversity, reads approach showed a much weaker correlation with assemblies approach ( $R^2$ = 0.3, 3083 genes). However, this result was for the nucleotide diversity of all those genes for which both values were available. When we restricted this comparison to our genes of interest, TFs and their target genes, this correlation improved substantially (Figure V2) ( $R^2$ = 0.64, 1237 genes). Our reads approach also underestimated nucleotide diversity. This was largely due to the difference in how gapped sites are treated in the MSA and in our approach. In an MSA, all columns with gaps are removed and there is information available for all sites of a gene. However, in our approach, the information is available only for sites which have a variant. As a result, the gapped sites with no variants are included in the calculation which leads to smaller values of nucleotide diversity. The alternative approach was to exclude the runs with any gaps but that led to even weaker correlation and so we decided to proceed with the former approach. With these limitations, it was necessary for us to ascertain if we can test our hypothesis using the reads approach. Therefore, we tested for the difference between non-synonymous diversity of TFs and corresponding averages of their TUs using both approaches and we found that both of these approaches supported the same conclusion that non-synonymous diversity of TFs was lower than their target genes. In fact, the assembly approach had stronger evidence toward the same conclusion (Figure V3).

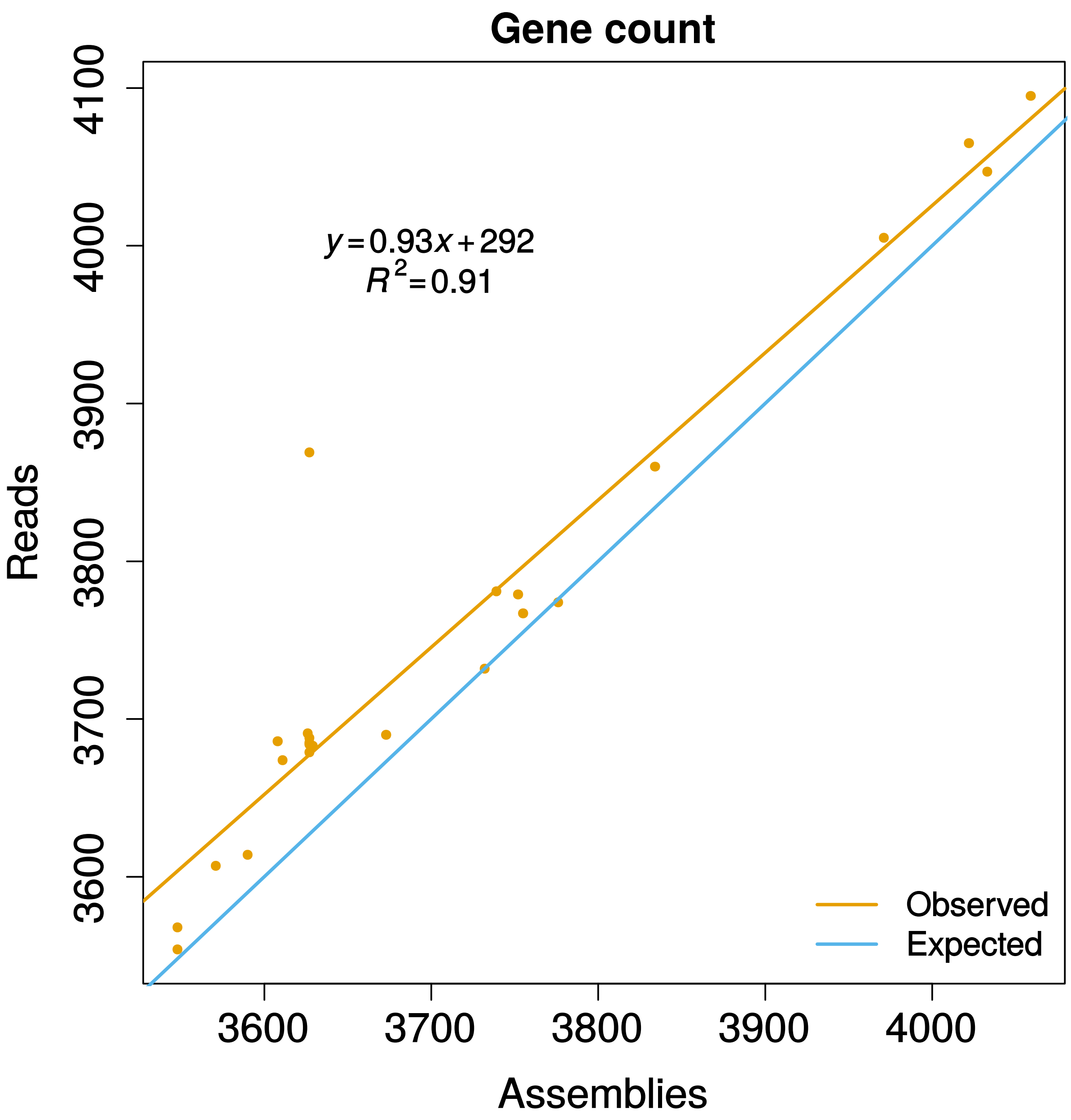


**Figure V1. Comparison of gene counts between assemblies and reads approaches.** Reads approach reliably infers gene presence, albeit with a slight overestimation of total count. This validation was based on 24 strains for which both reads data and assembled sequence was available.

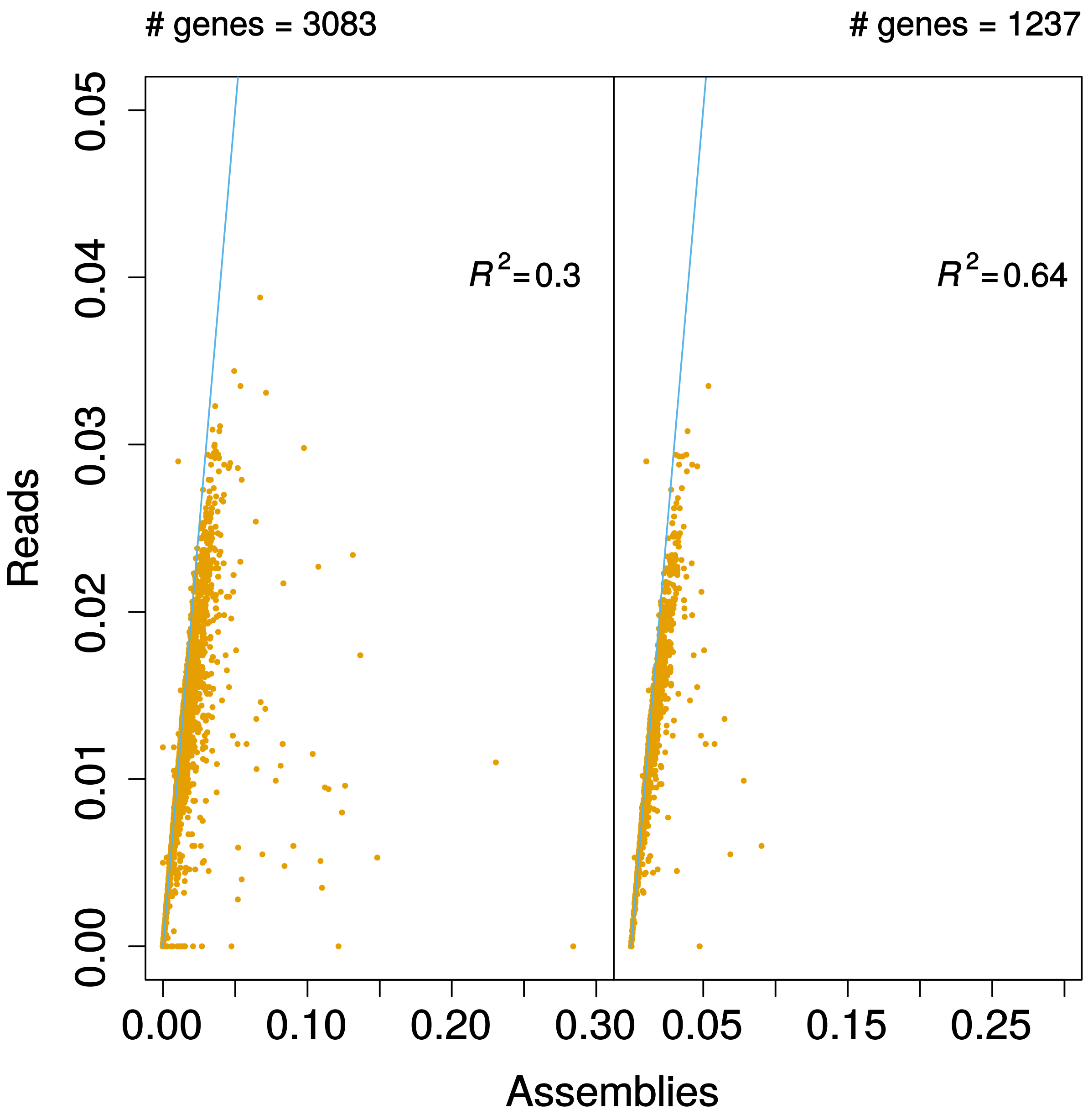


**Figure V2. Comparison of nucleotide diversity between reads and assemblies approaches.** Reads approach was a poor predictor of nucleotide diversity when a large number of *E coli* genes were analyzed. However, the correlation, as estimated by the Pearson method, improved substantially upon limiting this comparison to genes of our interest *i.e.*, TFs and their TGs. Reads approach mostly underestimated diversity. See supporting text for an explanation.

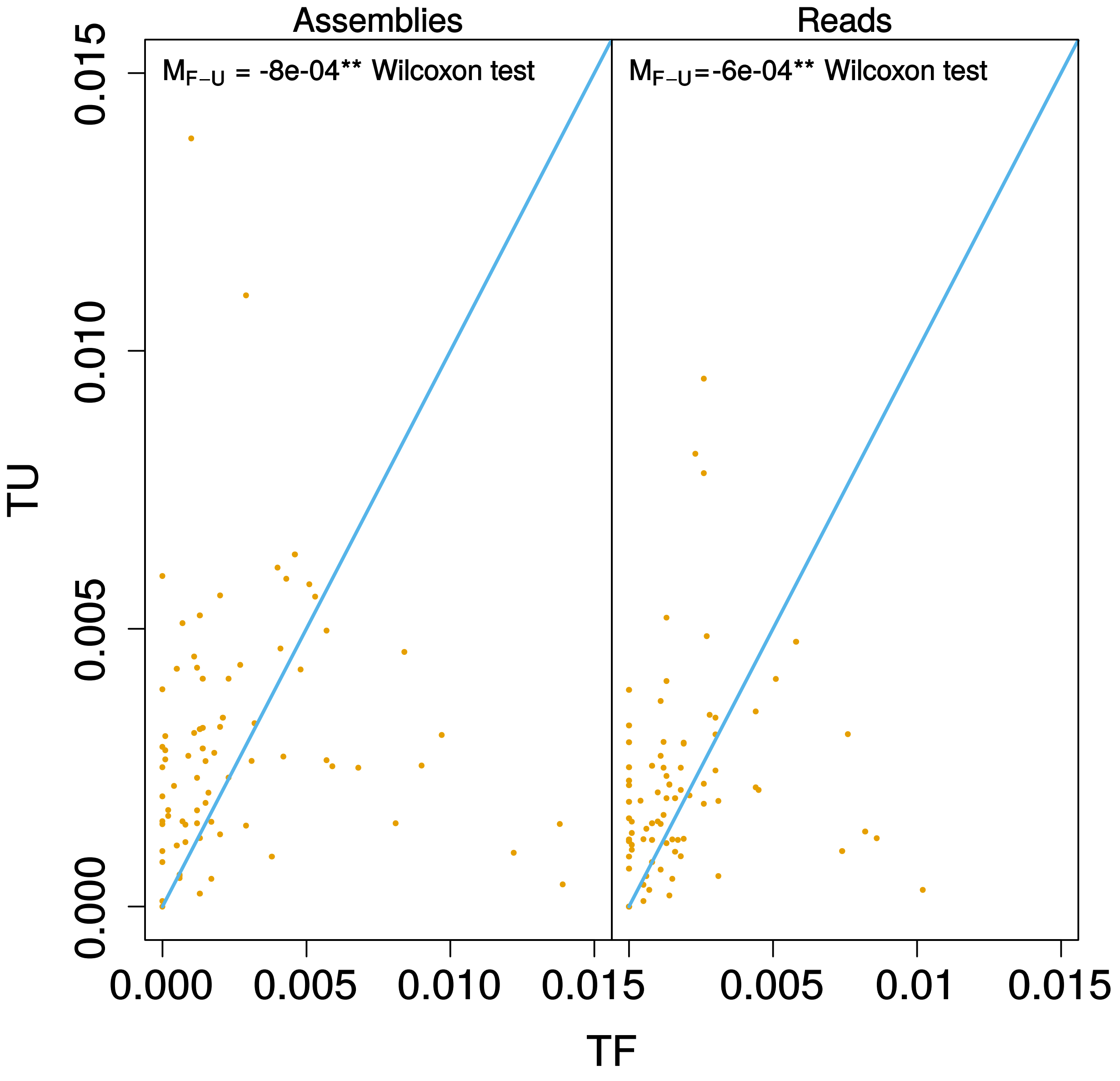


**Figure V3. Comparison of differences in non-synonymous diversity of TFs and their TGs between reads and assemblies approaches.** Using reads approach, we concluded that TFs have less non-synonymous diversity than their TGs. It was also true for the validation set. The same conclusion was reached with the assemblies approach. If anything, the strength of evidence against the null hypothesis was more with assemblies approach. Wilcoxon signed rank test: $P_{reads} = 0.005$, $P_{assemblies} = 0.001$.
